# A Versatile Photocrosslinkable Silicone Composite for 3D Printing Applications

**DOI:** 10.1101/2023.07.17.549429

**Authors:** Mecit Altan Alioglu, Yasar Ozer Yilmaz, Ethan Michael Gerhard, Vaibhav Pal, Deepak Gupta, Syed Hasan Askari Rizvi, Ibrahim T. Ozbolat

## Abstract

Embedded printing has emerged as a valuable tool for fabricating complex structures and microfluidic devices. Currently, an ample of amount of research is going on to develop new materials to advance its capabilities and increase its potential applications. Here, we demonstrate a novel, transparent, 3D printable, photocrosslinkable, and tuneable silicone composite that can be utilized as a support bath or an extrudable ink for embedded printing. The proposed silicone composite can be tuned to achieve ideal rheological properties, such as optimal self-recovery and yield stress, for use in 3D printing. When used as a support bath, it facilitated the generation microfluidic devices with circular channels of diameter up to 30 µm. To demonstrate its utility, flow focusing microfluidic devices were fabricated for generation of Janus microrods, which can be easily modified for multitude of applications. When used as an extrudable ink, 3D printing of complex-shaped micro- and macro-constructs were achieved with integrated electronics, which greatly extends its potential applications towards developing complex flexible parts for soft robotics and prosthetics. Further, its biocompatibility was tested with multiple cell types to validate its applicability for medical and tissue engineering use. Altogether, this material offers a myriad of potential applications in material and medical fields by providing a facile approach to develop complicated 3D structures and interconnected channels that can further advance microfluidics and soft-robotics research.

## 1. Introduction

Three-dimensional (3D) printing has opened a plethora of opportunities in manufacturing soft matters^1–3^. Specifically, embedded 3D printing has recently gained substantial attention in developing complex-shaped functional biological constructs with soft materials, which were nearly incompatible with conventional 3D printing technologies. Embedded 3D printing is a method that involves depositing soft materials into support baths in a freeform manner to produce intricate 3D structures with specific shapes and architectures. It has been greatly explored in the field of tissue engineering, regenerative medicine, and biomedical research by printing biological materials, including cells, tissue spheroids, or acellular bioinks, within a scaffold or matrix that imitate the natural tissue environment^4–6^.

Embedded 3D printing with silicone offers promising potential in microfluidics research as silicone has distinctive properties and capabilities, such as biocompatibility without eliciting significant immune responses^7^, flexibility, elasticity, chemical inertness, durability, and optical transparency, which makes it an ideal material for the abovementioned application^8–11^. Further, silicone-based materials demonstrate excellent resistance to a broad spectrum of temperatures, chemicals, and mechanical forces, thus they can handle fluid flow under high pressures and temperatures. They also enable laminar flow patterns, accurate fluid mixing, and separation, thereby facilitating diverse chemical and biological processes^12^. Moreover, they offer single-cell analysis and diagnostics in precision medicine thus facilitating research on cellular heterogeneity, cancer detection, genetic screening, and personalized therapies. Embedded printing of silicone-based microfluidics has recently gained interest in the biofabrication and tissue engineering fields as well^13^.

Photocrosslinkable silicone is advantageous over its thermally crosslinkable counterparts due to its tunable crosslinking, spatial control, rapid and on-demand fabrication, compatibility with biological samples, integration with other materials, and chemical resistance^14^. It can facilitate the formation of microfluidic channels, chambers, and valves with accurate dimensions. Furthermore, photocrosslinking ensures thorough solidification of the material, thereby diminishing the risk of biological sample contamination or leaching. Photocrosslinked silicone-based microfluidic devices exhibit favorable characteristics for utilization in various fields, such as cell culture, tissue engineering, and biomedical research^15^.

In this work, we developed a transparent photocrosslinkable silicone composite (PhSC) that had tunable rheological and mechanical properties. The adaptability of PhSC was demonstrated by using it both as a support bath and an extrudable ink for embedded printing. When used as a support bath, microfluidic devices were fabricated without any need for complex fabrication processes and cleanroom facilities. To exemplify its use, microfluidic devices were utilized to generate microgels, including Janus microrods. When utilized as a printable ink, complex-shaped structures were fabricated, such as but not limited to a hand model with a pressure sensor to demonstrate its capabilities towards soft robotics or custom prosthetics. Lastly, the biocompatibility of PhSC was also demonstrated to validate its potential for future biological applications.

## 2. Materials and Methods

### 2.1. Materials

Vinyl terminated (4-6% diphenylsiloxane) dimethylsiloxane copolymer (PDV-0525) (vtS) and (mercaptopropyl 13-17%) methylsiloxane - dimethylsiloxane copolymer (SMS-142) (mcpS) were acquired from Gelest (Morrisville, PA, USA). HMDS-treated silica (AEROSIL R 812 S) (Si) was acquired from Evonik (Essen, Germany). Thixotropic agent THI-VEX™ (ThA) was obtained from Smooth-On (Macungie, PA, USA). 2,2-Dimethoxy-2-phenylacetophenone (DMPA), xanthan gum (XG) from Xanthomonas campestris, gelatin from cold water fish skin, light mineral oil, Span 80, and Pluronic F-127 were acquired from Sigma-Aldrich (St. Louis, MO, USA). Lithium phenyl (2,4,6-trimethylbenzoyl) phosphinate (LAP) acquired from TCI chemicals (Portland, OR, USA).

### 2.2. Support Bath Preparation

PhSC support baths were prepared by mixing vtS, mcpS, Si, ThA, and DMPA in a speed mixer DAC-330-110 SE (Flacktek Manufacturing, Landrum, SC, USA). Preparation of a 20 g PhSC that had 16% mcpS and 4% Si concentration was as follows: First, 15.2 g vtS (72% w/w) and 3.2 g mcpS (16% w/w) were placed in a 40 ml container and mixed at 3,500 rpm for 1 min. Then 0.8 g (4% w/w) Si was added to the container and mixed at 3,500 rpm for 5 min. Then 0.8 g (4% w/w) ThA was added to the container and mixed at 3,500 rpm for 2 min. Finally, 10% (w/v) DMPA was dissolved in ethanol and 40 µL (0.2 % w/v) was added to the container and mixed at 3,500 for 5 min. After mixing was completed, PhSC was loaded into 5 or 10 mL syringes and centrifuged at 4,000 rpm for 5 min to remove air bubbles. After centrifugation, syringes that contain PhSC were stored in dark at room temperature to prevent undesired photocrosslinking. After PhSC was cast, it was dipped in water bath to remove contact with the atmosphere to eliminate the effects on oxygen on photocrosslinking^16^. Then, PhSC was crosslinked by exposing 405 nm light for 10 min with a 358 mW/cm^2^, 405 nm light source (CR-UV light, Creality, Shenzhen, China).

### 2.3. Transparency Measurements

Transparency measurements were conducted according to our previous studies^13, 17^. Briefly, optical absorbances were measured using a spectrophotometer (BioTek, Winooski, VT, USA). To measure transparency, PhSC samples with 4, 8, 16, and 32% mcpS were prepared and loaded into a 96 well plate (100 µL per well). After loading, well plates were centrifuged at 1,600 rpm for 5 min. Then, single point measurements at 450 nm (blue), 560 nm (green), and 660 nm (red) wavelengths were conducted before and after photocrosslinking of PhSC. UV-Vis spectrum measurement from 340 to 990 nm was also conducted after photocrosslinking. To display transparency, PhSC samples were placed on a paper with printed incremental font sizes ranging from 2 to 10 points and photographed.

### 2.4. Mechanical Testing

The American Society for Testing and Materials (ASTM) standard D638-14 was used to prepare and measure samples for tensile tests. PhSC samples with 8, 16, and 32% mcpS were prepared and cast into molds with Type V dog-bone shapes. After casting and crosslinking, samples were tested on an Instron 5966 system (Instron, Norwood, MA, USA) with a 1 kN load cell and pneumatic tensile grips. Samples were subjected to tension at a rate of 10 mm/min until failure.

The ASTM standard D395-18 was used to prepare and measure samples for unconfined compression tests. PhSC samples with 8, 16 and 32% mcpS were prepared and cast into cylindrical molds with 6 x 13 mm (height x diameter) dimensions. After casting and crosslinking, samples were tested on the Instron 599 system with a 10 kN load cell. Samples subjected to compression at a rate of 1 mm/min to failure or to 90% strain.

The ASTM standard D2240-15 was used to prepare and measure samples for durometer testing. Tekcoplus (Hong Kong, China) shore A durometer was used to measure the shore A hardness of the samples, and the values were reported as Hardness A (HA).

To incorporate the Mooney-Rivlin material model, engineering stress-strain curves from the compression and tensile experiments were stitched together, the engineering strain 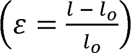 was converted to stretch ratio 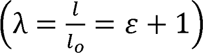 with *l* being the current length and *l*_0_ the initial length of the specimen. We used the machine generated engineering stress (σ) that was mathematically the principal component of the first Piola-Kirchhoff stress tensor in the direction of sample deformation. A formulation for the first Piola-Kirchhoff stress response under uniaxial extension or compression of the incompressible Mooney-Rivlin material was used^18, 19^.

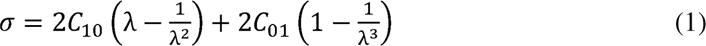

Here, *C_10_* and *C_01_* were empirically determined material constants for a Mooney-Rivlin hyper-elastic material. For the curve fitting, we used the curve-fitting toolbox from MATLAB. In the small strain regime, the shear modulus (μ) for incompressible material can be calculated from these material constants as:

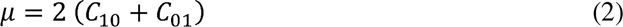

Elastic modulus can be calculated from the following formula.

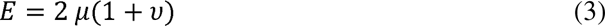

Where υ is Poisson’s ratio, which is approximately 0.5 for silicones since silicones are nearly incompressible materials^20^.

### 2.5. Nuclear Magnetic Resonance (NMR) Measurements

Proton NMR of polymers was assessed in deuterated acetone at a 5 mg/mL concentration using a Brucker NEO-400 (Billerica, MA, USA) at 500 MHz and room temperature with a total of 32 scans and analyzed using Brucker TopSpin 4.2.0 software.

### 2.6. Crosslink Density Measurements

Density (*p*) was assessed using the density accessory for the Mettler Toledo XP504 balance (Mettler Toledo, Columbus, MN, USA) at room temperature, where DI water was used as an auxiliary fluid. Briefly, 2 mm cubic samples were first weighed in air to determine (*W_a_*). Samples were then immersed in the auxiliary fluid bath to determine (*W_b_*). *p* (g/cm^3^) was then calculated according to equation (4)

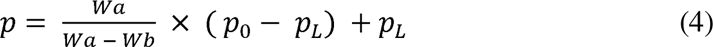

where *p*_0_ is the density of DI water, and *p_L_* is the density of air (0.0012 g/cm^3^). Crosslink density (*v*) and molecular weight between crosslinks (*M_C_*) were calculated according to equation (5):^21^

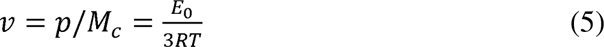

where *E_0_*is Young’s modulus, *R* is the universal gas constant (8.3144 J/mol K), and *T* is the absolute temperature (K).

### 2.7. Determination of Gel Fraction

Gel fraction (GF) was determined via extraction of remaining free oligomers from crosslinked organogels. Briefly, rectangular gel samples weighing approximately 60 mg were incubated in 1,4 Dioxane (10 mL per sample) under mild shaking at room temperature for 24 h followed by lyophilization for 4 days to remove the residual solvent. GF (%) was calculated according to equation (3):^22^

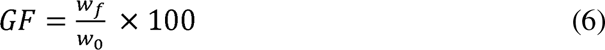

Where *w*_0_ and *w_f_* were original and final weights of samples, respectively.

### 2.8. Rheology Measurements

To conduct rheology experiments, PhSC samples with 3, 4, and 5% Si were loaded on a MCR 302 rheometer (Anton Paar, Graz, Austria) equipped with a 25 mm parallel plate. Measurements were conducted at room temperature. The amplitude sweep test was performed within a range of 0.1 to 200 Pa shear stress at a frequency of 1 Hz. The shear rate sweep test was performed within a range of 0.1 to 100 1/s shear rate. The frequency sweep test was performed within a range of 0.1 to 100 rad/s angular frequency. The thixotropic recovery sweep test was performed by applying 0.1 and 100 1/s shear rates with 60 s intervals.

### 2.9. Embedded Printing of the Sacrificial Ink

To develop a sacrificial ink, first, 3% (w/v) XG stock solution was prepared by blending it in DI water for 4 min in a commercial blender (Magic Bullet, Home Appliances, Los Angeles, CA, USA) and Pluronic F-127 30% (w/v) was prepared by mixing it in DI water using a magnetic stirrer at 4 °C overnight. Then, XG and Pluronic were mixed at a 1:1 ratio to obtain the sacrificial ink. After blending, sacrificial ink was loaded in a 3 mL syringe and centrifuged at 4,000 rpm for 5 min. Then, 3 mL syringe was loaded onto a 3D bioprinter (Inkcredible+, Gothenburg, Cellink, Sweden) for embedded printing. Printing characterization was performed by printing the sacrificial ink inside PhSC with a 34G (60 µm inner and 200 µm outer diameter) needle. The relationship between microchannel diameter and printing parameters was investigated by incrementally increasing the print speed from 15 to 120 mm/min at a 50, 60, 80 and 100 kPa printing pressure. For roundness measurements, Si microchannels were sliced with a blade to obtain cross-sections. These cross-sections were imaged with a Zeiss Axio Observer (Oberkochen, Germany) and analyzed using ImageJ (National Health Instruments (NIH), Bethesda, MA, USA).

### 2.10. Fabrication of Microfluidic Devices

Microfluidic devices were fabricated by printing the sacrificial ink in the PhSC support bath with the Inkcredible+ bioprinter. To achieve this, PhSC was prepared and cast into a 3D printed polylactic acid or laser-cut acrylic container. Then, the sacrificial ink was loaded into the bioprinter and 3D printed inside the PhSC support bath. G-codes for printing were generated using MATLAB (MathWorks, Natick, MA, USA). After printing was completed, PhSC was placed inside a water bath and photocrosslinked using the 405 nm light source. Then the sacrificial ink was removed from microfluidic devices by perfusing water through channels. Removal of sacrificial ink was done effortlessly since sacrificial ink was water based and flushed out easily. To achieve flow inside the microfluidic devices, syringe pumps (New Era Pump System Inc., Farmingdale, NY, USA) were utilized.

### 2.11. Generation of Microgels

Microgels, homogenous microrods and Janus microrods, were produced using flow-focusing microfluidic devices. Flow-focusing microfluidic devices were fabricated via embedded printing in PhSC. Methacrylated fish gelatin (GelMA) was synthesized using a previously reported method^23^. To obtain aqueous phase, 15% (w/v) GelMA and 5% (w/v) LAP was dissolved in DI water. To obtain oil phase, 97% (v/v) light mineral oil and 3% (v/v) Span 80 was mixed using the speedmixer. Oil and aqueous phase were injected in the flow focusing device with syringe pumps to obtain microgels. Oil flow was kept at 20 µL/min and GelMA flow was increased from 1 to 20 µL/min to investigate the effect of different flow speeds on microgel formation. Microgels were imaged with an Evos XL Core microscope (Life Technologies, Carlsbad, CA, USA) and their size was quantified using ImageJ.

### 2.12. Embedded Printing of the PhSC Ink in a Sacrificial Support Bath

To achieve embedded printing of the PhSC ink, two different sacrificial support baths were tested. XG sacrificial bath was prepared by blending 2% (w/v) XG in DI water with a commercial blender for 4 min. Mineral oil based sacrificial support bath was prepared by mixing 8% (w/v) of silica filler in light mineral oil using the speedmixer at 3,500 rpm for 5 mins. After sacrificial support baths were prepared, PhSC was prepared and loaded into the bioprinter. PhSC was printed inside the sacrificial support bath using a 25G (250 µm inner and 520 µm outer diameter) needle. G-codes were generated using MATLAB and Cura (Ultimaker, Ultrecht, Netherlands). Next, the 3D printed constructs and devices were photocrosslinked. Then the support bath was removed, and the constructs and devices were stored for future use.

### 2.13. Cell Culture

Commercially available human adipose derived stem cells (ADSCs) (Catalog number: PT-5006, Lonza, Walkersville, MD, USA) were cultured in HyClone Dulbecco’s modified Eagle medium (F12) (Hyclone, Marlborough, MA, USA) and expanded from passage 1–3 to for experiments. The media was supplemented with 1% U ml^−1^ Penicillin/Streptomycin (Corning, Manassas, VA, USA) and 20% fetal bovine serum (R&D Systems, Minneapolis, MN, USA). The cells were cultured in incubator maintained at 37 °C and 5% CO_2_ and the cell media was changed on alternate days. Human dermal fibroblasts (HDFs), obtained from N. Zavazava’s lab at the University of Iowa (Iowa City, IA, USA), were cultured in DMEM (Corning) supplemented with 1% glutamine (Thermo Fischer Scientific, Waltham, MA, USA), 10% fetal bovine serum (Corning), 1% Penicillin-Streptomycin (Corning), and 1% sodium pyruvate (Thermo Fischer Scientific).

To measure cellular activity, PhSC was cast into 24-well plates, centrifuged, crosslinked, and sterilized with 70% ethanol overnight. For each well, 5 × 10^4^ HDFs and ADSCs were seeded on PhSC that was coated with 0.1% gelatin (Gelatin from porcine skin, G2500, Sigma-Aldrich). Cell viability was measured at Days 1, 4, and 7. For the viability assessment, ADSCs on the PhSC surface were rinsed with Dulbecco’s phosphate-buffered saline (DPBS) (Corning) three times and stained using calcein AM (2 µl/ml) and ethidium homodimer-1 (EthD-1) (2 µl/ml) (Thermo Fischer Scientific). Then plates were incubated with staining solutions for 1 h in an incubator. After incubation, wells were rinsed with DPBS three times and imaged using the Zeiss Axio Observer. The cell viability was quantified by analysing the obtained images using ImageJ (NIH).

To evaluate cell proliferation, for each well, 5 × 10^4^ ADSCs were seeded on 0.1% gelatin-coated PhSC. Proliferation was evaluated at Days 1, 4, and 7. Alamar blue dye reduction assay (Invitrogen, Waltham, MA, USA) was performed as per manufacturer’s protocol. Briefly, the seeded cells were incubated with 400 µL of 10% (v/v) of the dye for 2 h at 37 °C. After incubation, fluorescence intensity at 570/590 nm (excitation/emission) was measured with a microplate reader (Tecan Infinite 200 Pro, Männedorf, Switzerland). The results were presented as percent of dye reduced, which were proportional to the number of viable cells present.

### 2.14. Statistical Analysis

All data were presented as mean ± standard deviation. Data were analyzed by GraphPad Prism and statistical differences were determined using one-way analysis of variance (ANOVA) with Tukey’s post hoc test, and the analysis, if fulfilling the null hypothesis at *p* ≤ 0.05 (*), *p* ≤ 0.01 (**) and *p* ≤ 0.001 (***), were considered statistically significant.

## 3. Results and Discussion

### 3.1. Development of PhSC Material for Embedded Printing of Microfluidics Devices

In this study, we developed a novel silicone composite (PhSC) that was photocrosslinkable. PhSC had yield-stress and self-recovering properties so that it was successfully used as a support bath for embedded printing of microfluidic devices. To fabricate microfluidic devices, first, PhSC was prepared by mixing vtS, mcpS, silica filler, thixotropic agent, and photoinitiator. After preparation, PhSC was cast into a container and the sacrificial ink was printed into PhSC. Then, PhSC was dipped in water and photocrosslinked using a 405 nm light source. (**Figure 1**) Although photocrosslinking occurred within a minute, PhSC was exposed to light for 10 min to ensure fully crosslinking. In the literature, one of the challenges of silicone-based microfluidic device fabrication is the need for degassing after casting^24^. However, this problem was eliminated with PhSC because reaction occurred rapidly without a thermal change and without bubbles generation, and thus the degassing step after casting was not required.

**Figure 1.**
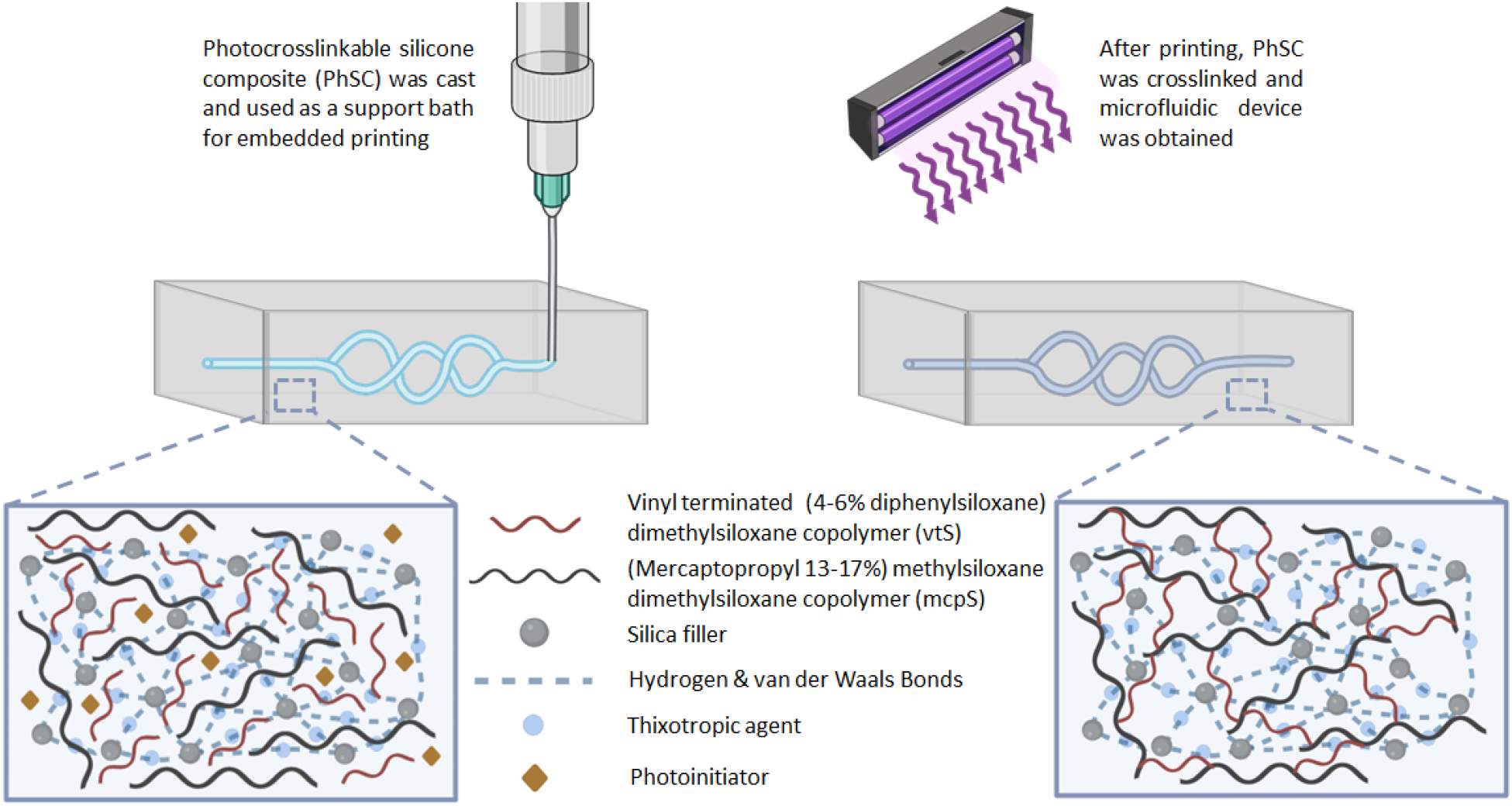
A schematic illustrating the development and utilization of PhSC as a support bath. Preparation, application, and contents of PhSC (created with BioRender.com).

To achieve photocrosslinking, thiol-ene click chemistry was chosen since it is a relatively simple process with high yields and reaction rates^25, 26^. A silicone polymer with vinyl ends and a silicone polymer that contains thiol (mercapto) group was used with a photoinitiator (PI). PI broke down to radicals with excitation from a 405 nm light source and photocrosslinking was obtained (**Figure 2**).

**Figure 2.**
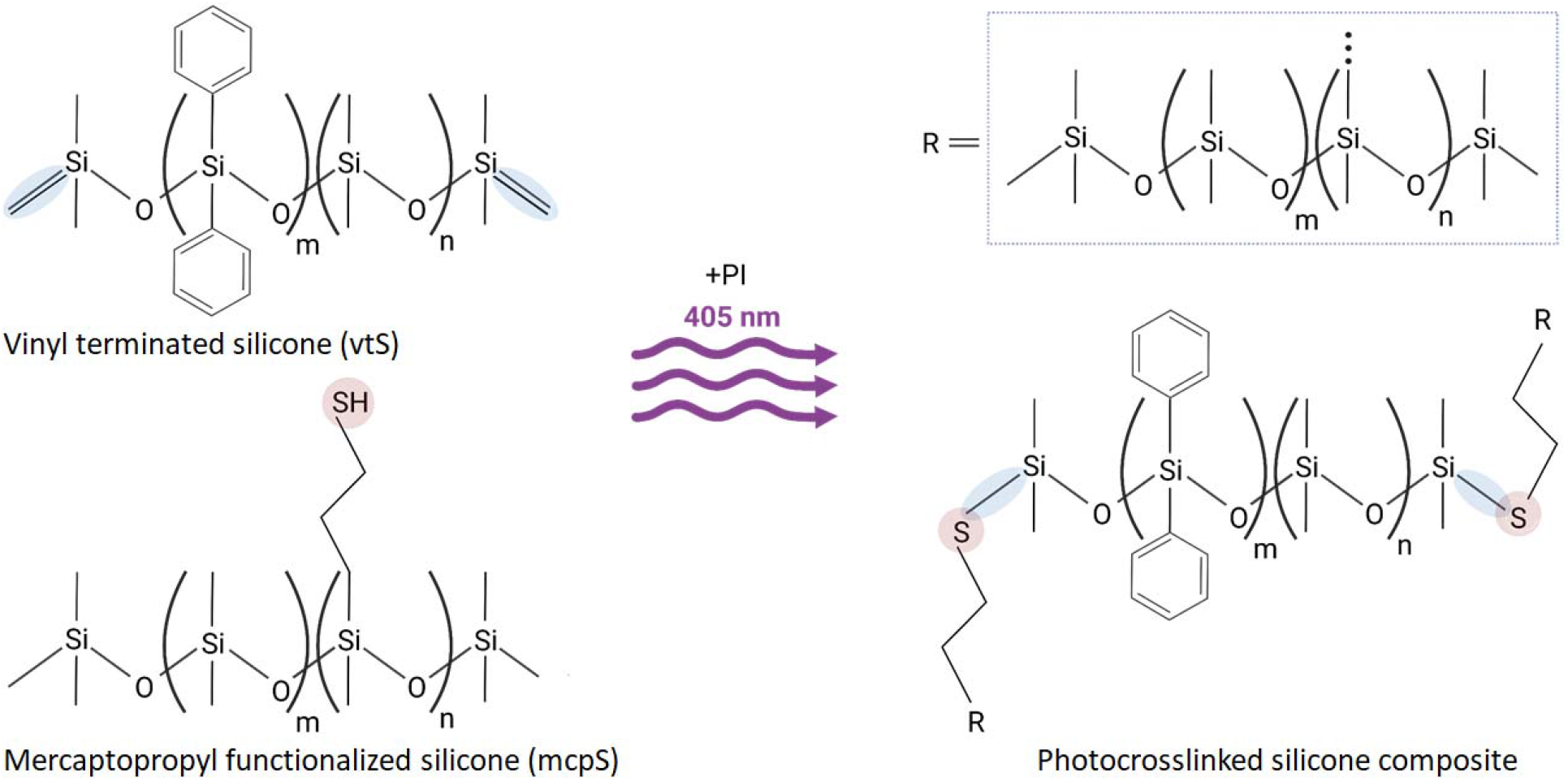
A schematic illustrating crosslinking of PhSC. Photocrosslinking of vinyl terminated silicone (vtS) and mercaptopropyl functionalized silicone (mcpS) by thiol-ene reaction.

### 3.2. Material Characterization

Optical transparency is a crucial property for the fabrication of microfluidic devices since microfluidic channels often are monitored during their target applications^9^. Also, optical transparency is additionally important for photocrosslinkable materials since light may get absorbed completely at the top regions of an opaque material and leaving the bottom regions of the material uncrosslinked. While preparing a silicone composite, optical clarity may be lost due to a number of factors, such as phase separation, optical reflectance mismatch, or inclusion of opaque additives. Here, we obtained a photocrosslinkable yield-stress silicone organogel that was highly transparent. Transparency of different compositions of PhSC was displayed in **Figure S1A.** As the concentration of mcpS changed from 4 to 32%, the transparency of PhSC did not alter significantly (**Figures S1B-E**). In addition, the crosslinking process did not affect the transparency considerably.

To characterize mechanical properties, tension and compression tests were performed as presented in **Figures 3A1, B1**. In our preliminary efforts, it was noticed that 4% mcpS was too soft to be loaded onto the pneumatic grips of the mechanical testing system; therefore, 4% mcpS concentration was excluded from the rest of the study. Mechanical properties of PhSC were highly tunable by changing the ratios of vtS and mcpS. 16% mcpS had the highest rigidity, followed by 8% mcpS. 32% mcpS samples had the highest strain before breaking and displayed considerable elasticity (**Figure 3A2**). The peak stresses ranged from ∼150 to 260 kPa as shown in **Figure 3A3**. The stress-strain curve of compression test also mirrored the findings of tension tests (**Figure 3B2**). Tensile peak stresses ranged from ∼3 to 20 MPa (**Figure 3B3**). The results of Shore A hardness test of PhSC samples were shown in **Figure 3C**. Hardness values were in the range of ∼1 to ∼17 HA. As we increased mcpS concentration from 8 to 16%, hardness increased; however, for the 32% group, a sharp decrease in hardness was observed.

**Figure 3.**
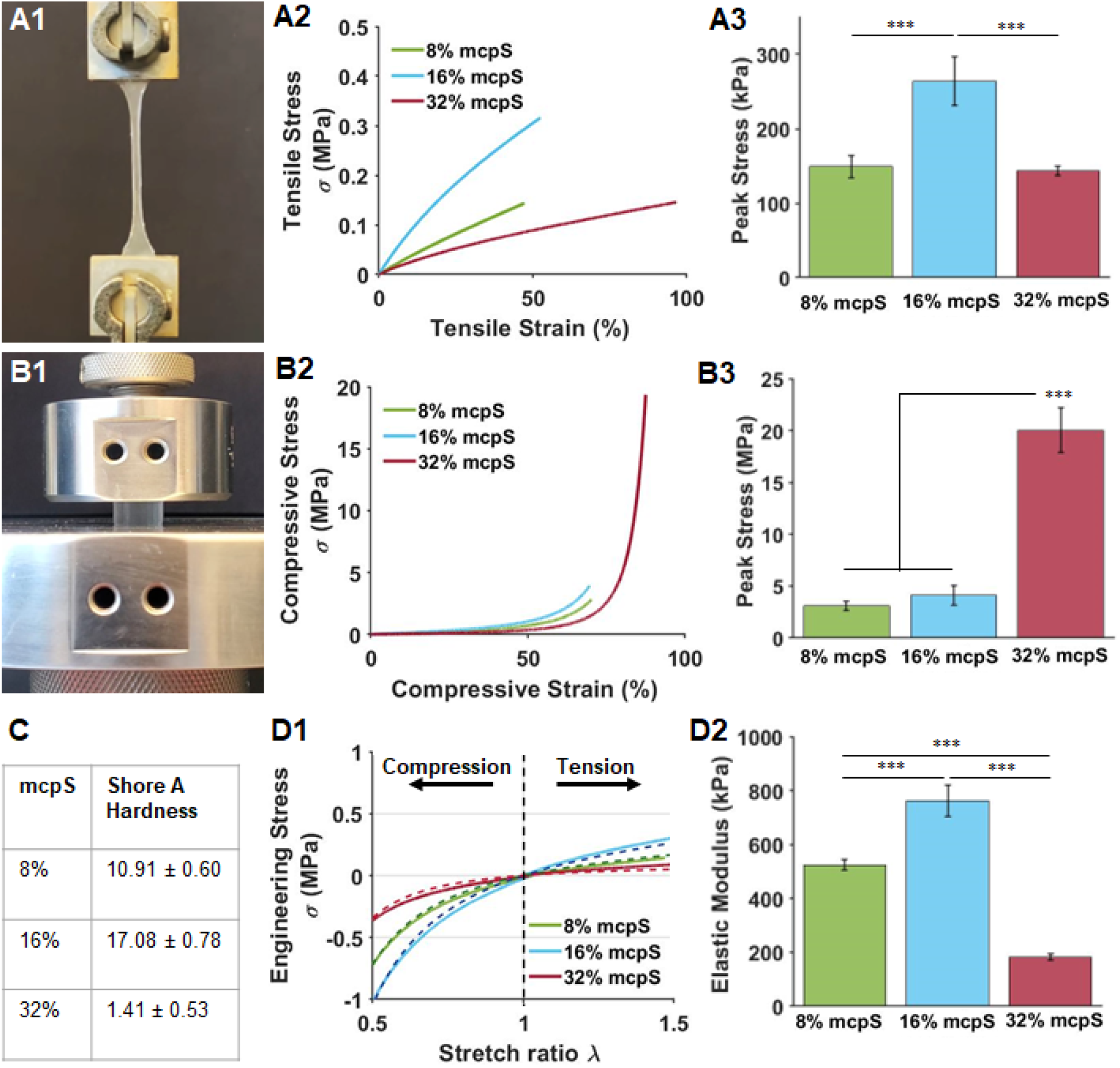
Mechanical characterization of PhSC. (A1) Tensile testing, (A2) tensile stress-strain curve, and (A3) tensile peak stress. (B1) Compressive testing, (B2) compressive stress-strain curve, and (B3) compressive peak stress. (C) Shore A hardness of PhSC samples. (D1) Mooney-Rivlin curve fitting on stress data. (D2) Elastic moduli of PhSC samples (*n*=4, *p*\*\*\*<0.001).

The developed PhSC displayed a non-linear elastic behavior. Small strain elasticity outcomes, such as elastic and shear modulus, are only applicable when strain is <1%. To have a complete description of the material, a non-linear material model should be incorporated. For that reason, one has hyper-elastic material models that provide a mathematical insight into material non-linearities. A non-linear isotropic and incompressible hyper-elastic material can store strain energy that can be classified by two independent strain invariants. We chose a two-parameter Mooney-Rivlin hyper-elasticity material model to analyze the hyper-elastic PhSC. Behavior of an incompressible Mooney-Rivlin material could be modeled with same parameters for both compression and tension of the material (**Figure 3D1**). The modulus of elasticity was significantly different for each sample (∼525 kPa for 8%, ∼760 kPa for 16% and ∼180 kPa for 32% mcpS) (**Figure 3D2**).

Network analysis of the obtained organogels was also conducted to describe material properties in details as tested above. Firstly, H-NMR analysis of the constituent polymers was performed, confirming the chemical structure of both vtS and mcpS (**Figure 4A**). Next, relative vinyl and thiol content was calculated based upon chemical structures and the molecular weight of the polymers. In brief, vtS contained two mols of vinyl group/mol for 142 µmol/g of polymer (Mn of vtS: 14,000 g/mol) while mcpS contains 0.15 mol thiol group/mol for 42.86 µmol/g of polymer (Mn of mcpS: 3,500 g/mol). Utilizing the above calculations and taking into account the relative weight percentages of each polymer, vinyl/thiol molar ratios of 34.99, 15.83, and 6.25 were obtained for 8, 16, and 32% mcpS, respectively. Considering the 1:1 nature of the thiol-ene click reaction, ratio of reactive groups could have a significant effect on network structure and resulting material properties, as observed via mechanical analysis (**Figure 3**), with an intermediate ratio of functional groups resulting in the stiffest material (16% mcpS) with maximal tensile properties (**Figure 3D2**). In contrast, more extreme ratios resulted in relatively soft, low modulus materials; however, it should be noted that compression properties were maximized in the latter case (32% mcpS, **Figure 3B3**). Further, 16% mcpS exhibited the highest level of crosslink density (**Figure 4B**). Given that material bulk densities were equivalent for all tested groups, a direct correlation between network structure and tensile properties was evident. Finally, gel fraction was found to be the greatest for 16% mcpS (**Figure 4C**), indicating the most complete reaction of polymers occurred within that composition. The network structure thus demonstrated a significant effect on material properties while allowing material tunability to match multiple potential applications. The presence of excess vinyl groups in all formulations could enable further material functionalization via click reactions performed simultaneous to or post gelation, particularly but not limited to functionalization with biological peptides or fluorescent tags, thus potentiating further applications.

**Figure 4.**
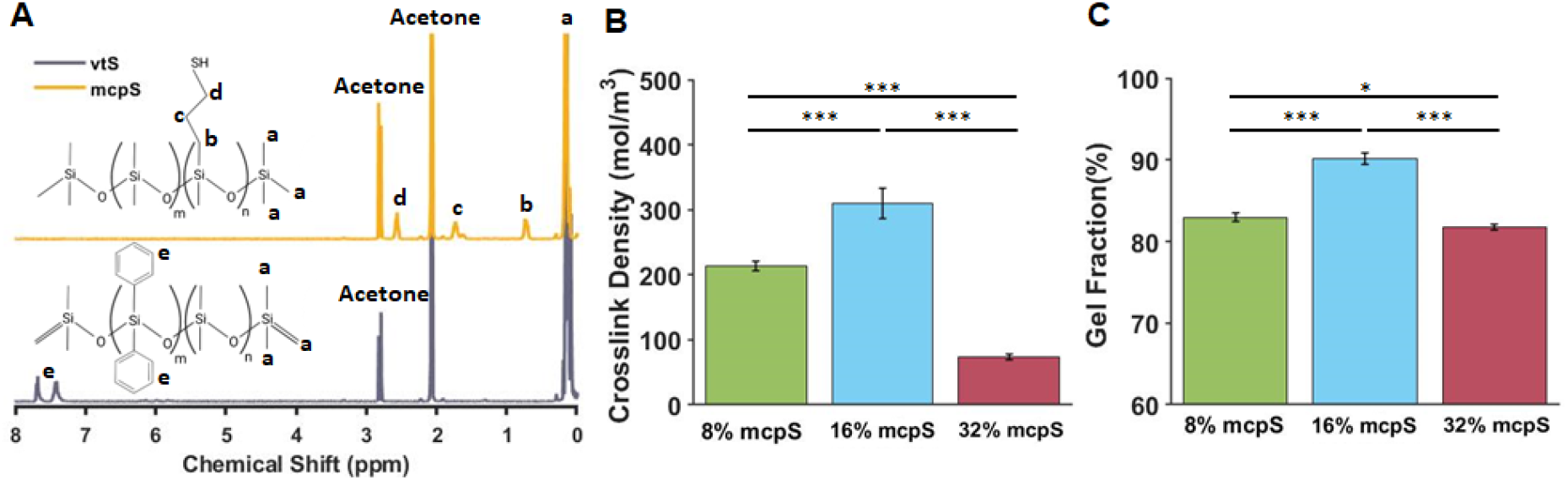
Analysis of Silicone Network. (A) H-NMR of vtS and mcpS mixed in deuterated acetone. (B) Crosslink density and (C) gel fraction of PhSC samples (*n*=4, *p*\*<0.05, *p*\*\*\*<0.001).

Rheological characteristics of PhSC were also examined to determine whether it was suitable to use as a support bath and a 3D printable ink. Our preliminary research demonstrated that the yield stress of 2% silica was too low, resulting in the displacement of the printed filaments during deposition, whereas the yield stress of 6% silica was too high, making it challenging to handle and cast PhSC. Therefore, a Si concentration range of 3 to 5% was examined for the rheology study. The viscoelastic behavior of PhSC was illustrated by the amplitude sweep test, as shown in **Figure 5A**. The outcomes showed a linear viscoelastic area, where storage modulus (G’) was greater than loss modulus (G’’) suggesting the material was in the gel state. The flow point, at which G’ was equal to G’’ suggested that the material changed from gel to liquid state, and finally the liquid state where G’ became greater than G’’. Since the viscosity of samples reduced as the shear rate increased, all of the formulations under investigation exhibited shear thinning behavior (**Figure 5B**). To model this viscoelastic silicone material, the Herschel-Bulkley model was used as below:^27, 28^

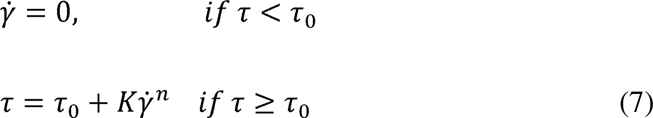

where 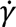 was the shear rate, τ was the shear stress, τ_0_ was the yield stress, *K* was the consistency index and *n* was the flow index. As the concentration of Si increased, yield stress of the mixture increased (**Figures 5C-D**). Frequency sweep tests were conducted within the linear viscoelastic region to investigate the Si-Co’s frequency dependence. Over the full range, all samples showed G’ > G’’ indicating the stable gel nature (**Figure 5E**). Finally, the self-recovering capability of PhSC was investigated with a 3-interval thixotropy test. As a high shear rate was applied and removed, the viscosity of the mixtures got completely recovered within seconds (**Figure 5F**).

**Figure 5.**
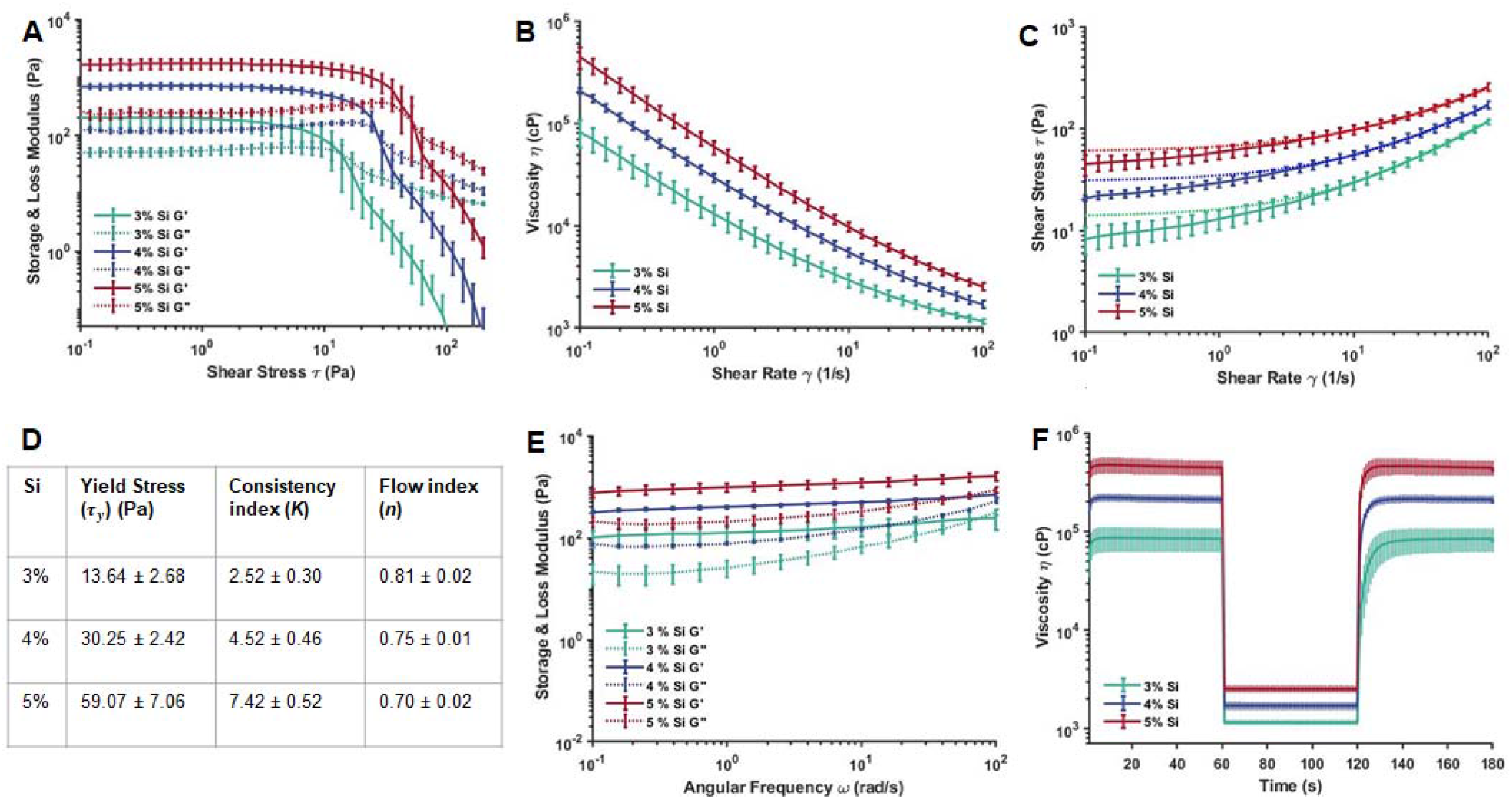
Rheological properties of PhSC. (A) Amplitude sweep test at a shear stress ranging from 0.1 to 200 Pa. (B) Shear rate sweep test at a shear rate ranging from 0.1 to 100 1/s. (C) Shear stress vs. shear rate curves, where dashed lines represent Hershel-Bulkley curve fittings on the shear stress data. (D) Frequency sweep test at a shear rate ranging from 0.1 to 100 rad/s. (E) 3-interval thixotropy test. The viscosity was measured at three intervals at alternating shear rates of 0.1 1/s for 60 s and 100 1/s for 60 s (*n*=6).

One of the significant advantages of photocrosslinkable silicone over thermally crosslinking silicone is their extended working life and pot life. Thermal crosslinking silicones are usually time dependent and once the final mixture is prepared, the user has a limited time to use the material before it crosslinks and solidifies. On the contrary, once the photocrosslinkable silicones are prepared, they can be stored in dark for long without initiating the crosslinking thus retaining their properties. To validate this, PhSC with 4% Si was prepared and kept in the dark for a week, and the rheological properties were tested. There was no change identified in the rheological characteristics of the stored samples as compared to the freshly prepared samples, and the final mixture was still crosslinkable and usable after a week (**Figures S2A-B**).

### 3.3. Embedded Printing in PhSC Support Bath

In our study, embedded printing was achieved by printing a sacrificial ink inside a yield-stress and self-recovering support bath (**Figure 6A**). We utilized PhSC as a support bath for embedded printing of microfluidic channels. Initially, XG was used as a sacrificial ink to achieve embedded printing. However, since XG ink was water based and PhSC was a hydrophobic material, the interfacial forces between the two materials were exerted to the printed filaments, which limited the minimum printable size of channels. To reduce the interfacial forces, XG was mixed with Pluronic F127. Since Pluronic had both hydrophilic and hydrophobic groups, it acted as an emulsifier and stabilized the printed filaments. With the addition of Pluronic, we were able to fabricate 30-µm straight channels (**Figures S3A-B**). However, <150 µm channels can only be printed along straight paths since it was challenging to print channels along paths with corners or branching networks.

**Figure 6.**
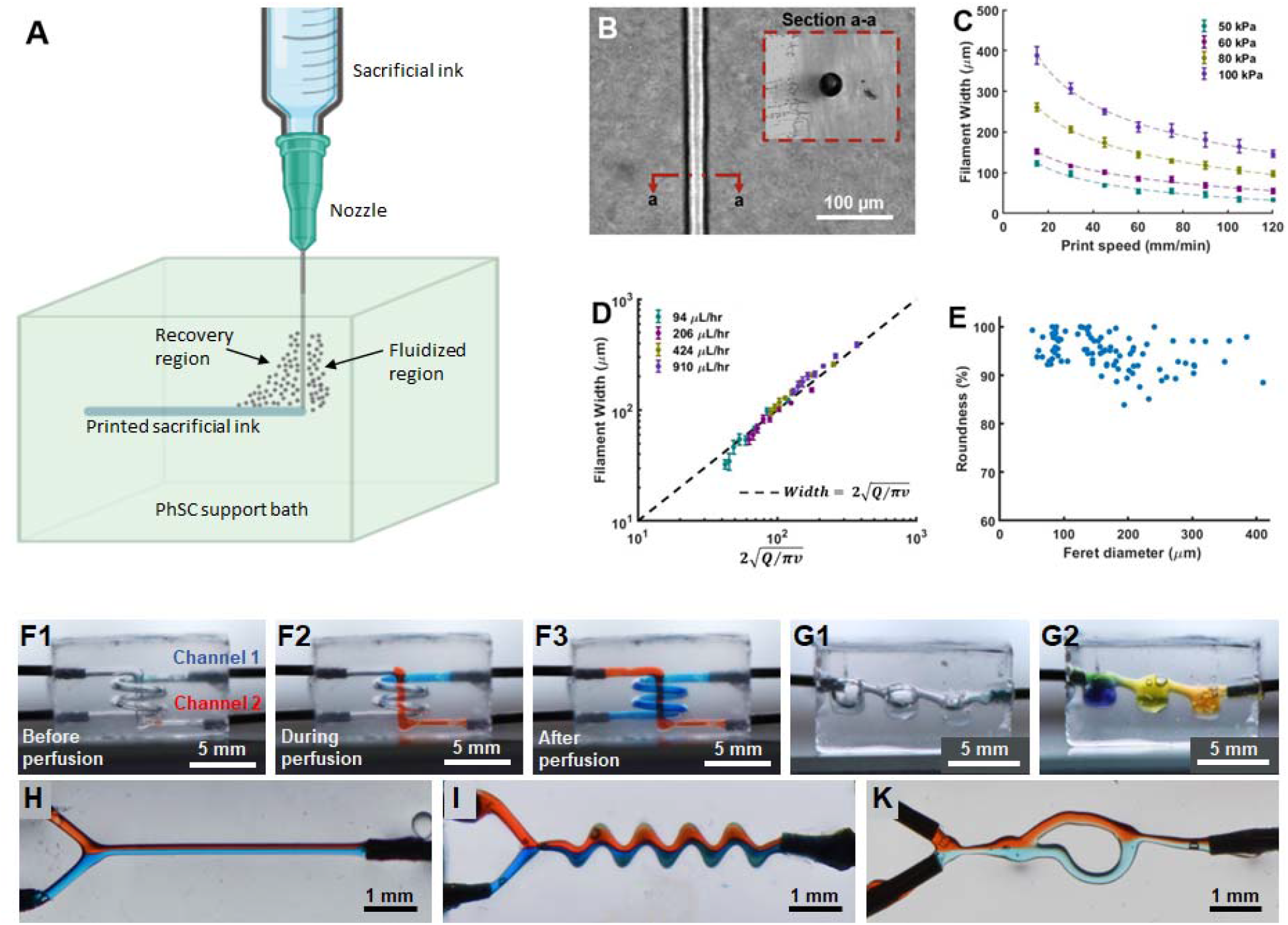
Embedded printing of the sacrificial ink inside PhSC. (A) A schematic of embedded printing. (B) A printed channel with a 30-µm diameter. (C) Filament width of microfluidic channels in line with fluid continuity equation. (D) Filament width versus print speed curves at 50, 60, 80 and 100 kPa printing pressures. (E) Roundness of printed microfluidic channels. 3D Printed (F) intertwined spiral and straight channel and (G) hydrogel chambers. Laminar flow in a (H) straight, (I) zigzag and (K) tesla-type valve channels.

The effects of printing parameters, such as extrusion pressure and printing speed, on the width and roundness of fabricated microfluidic channels were examined. 34G (60 µm inner and 200 µm outer diameter) blunt straight needles were used for the extrusion of sacrificial inks. Conical needles were not chosen because they would produce a greater yield zone around the needle since their outer diameter increases with distance from the nozzle tip. A 30-µm channel with a circular cross-section was shown in **Figure 6B**. Microchannel sizes were controllable by changing the printing parameters. As the printing speed was increased, the microchannel width was reduced and while the extrusion pressure increased, the microchannel width increased (**Figure 6C**). Also, width of microchannels was in line with fluid continuity equation (**Figure 6D**). **Figure 6E** shows the roundness of the channels and it was observed that the channels were highly circular. To demonstrate the printing capabilities, a complex shape having a vertical straight channel with an enclosing spiral channel was fabricated (**Figure 6F**). This validates that embedded printing in PhSC could be used to 3D print complex structures and channels to achieve intricate microfluidic devices. These devices could have chambers to house different materials. To prove this, chambers were 3D printed inside PhSC and loaded with hydrogels with different pH values. A universal pH indicator was perfused through the device to visualize hydrogels’ pH differences (**Figure 6G**). Also, laminar flow in channels was demonstrated by flowing dyed liquids in a straight (**Figure 6H**), zigzag (**Figure 6I**) and tesla-type valve^29^ (**Figure 6K**).

### 3.4. Flow-Focusing Microfluidic Device Fabrication

To demonstrate the utility of embedded printing, flow-focusing microfluidic devices were designed and printed manufactured. These devices were used to generate microgels by flowing GelMA and mineral oil though different channels. When GelMA aqueous phase and oil phase merged on the flow focusing junction, they separated and formed microgels. To produce microrods, devices were designed with extended channels (**Figure 7A**) to achieve UV crosslinking of microgels before they were collected at the outlet, similar to published research^30^. After microgels were formed at the flow-focusing region, they might travel through the channels in a cylindrical chape (**Figure 7B**, **Movie 1**) and when they were crosslinked at this later stage, they formed microrods. Further, the effect of flow rate on microgel size and shape was investigated (**Figures 7C-D**). The oil flow speed was kept at 20 µL/min and GelMA flow speed was increased from 1 to 20 µL/min. At 1 µL/min, only spherical microgels with a diameter ranging from 30 to 160 µm were formed. As the flow speed was increased to 10 µL/min, microgels exhibited elongated shapes. At 20 µL/min, we obtained microrods with lengths reaching up to 700 µm. We also modified the design to achieve Janus microrods (**Figure 7H**). In this design, two aqueous phases were first merged and formed a continuous non-mixing two phase flow. This two-phase flow merged with the oil phase at the flow-focusing junction to form Janus microrods (**Figure 7I**, **Movie 2**). To the best of our knowledge this was the first demonstration of Janus microrods produced with a flow-focusing device. These Janus microrods could be utilized for applications where two or more types of merged hydrogels are needed.

**Figure 7.**
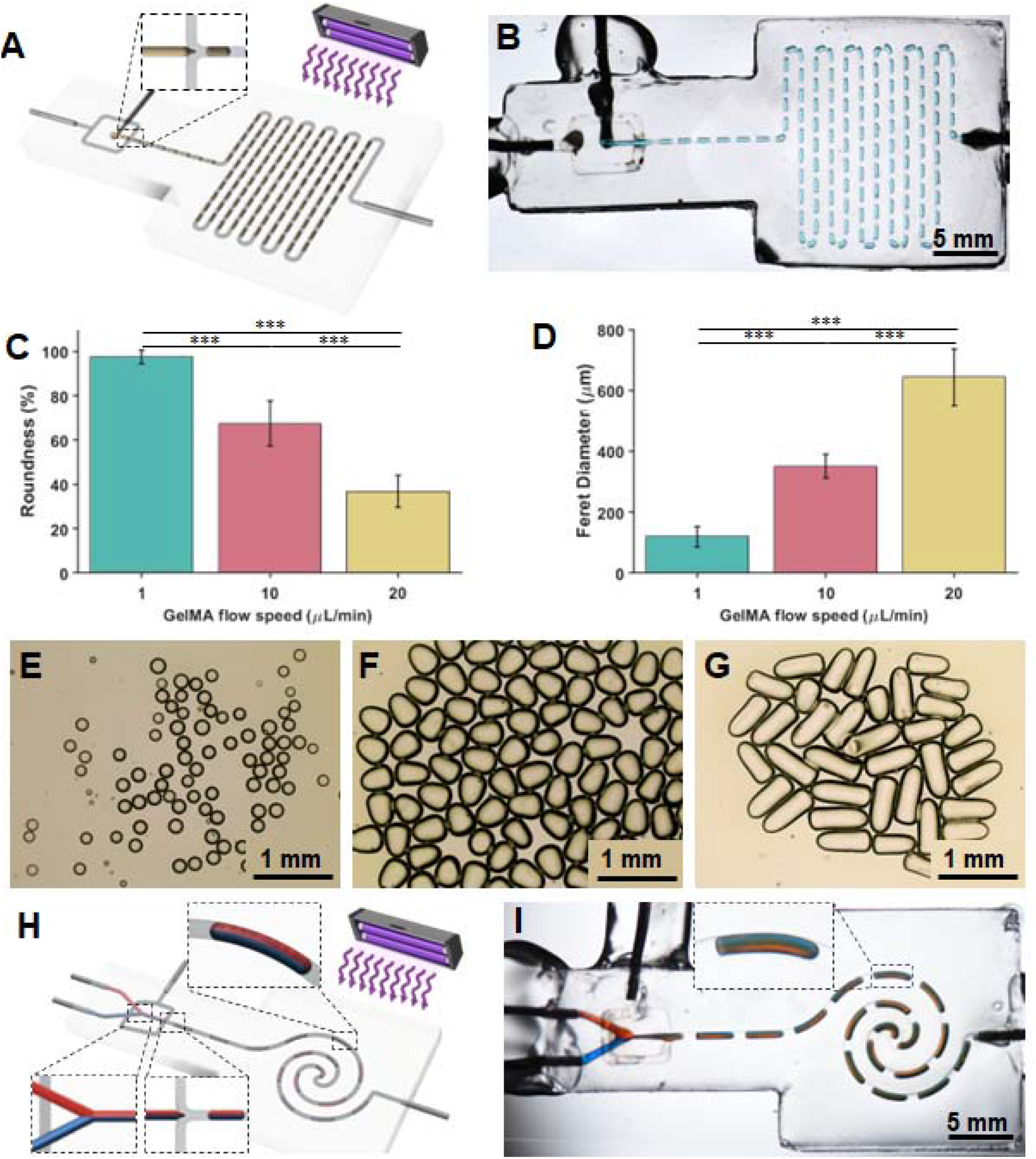
Flow-focusing microfluidic devices. (A) A schematic of flow focusing microfluidic device design and application. (B) A flow-focusing microfluidic device generating microrods. (C) Roundness of produced microgels with different GelMA flow speeds. (D) Feret diameter (length) of produced microgels with different GelMA flow speeds. Microgels produced with (E) 1, (F) 10, and (G) 20 µL/min GelMA flow speed. (H) A schematic of flow-focusing microfluidic device design and application for Janus microrods. (I) Flow-focusing microfluidic device generating Janus microrods.

### 3.5. PhSC as an Extrudable Ink

PhSC is versatile and can be used as a 3D printable ink as well. To achieve printing, PhSC with 4% Si was loaded into a bioprinter and printed inside a sacrificial support bath. After printing, printed structures were photocrosslinked and removed from the sacrificial support bath (**Figure 8A**). Initially both hydrophilic XG support bath and hydrophobic mineral oil support bath were tested. Although the hydrophobic support bath enabled embedded printing and had decent printability, PhSC ink interacted with the support bath and turned opaque (**Figure S4**). Therefore, for the following prints, XG support bath was utilized. To demonstrate printing capabilities, a bacteriophage model (**Figure 8B**) and tibia model (**Figure 8C**) were printed. Tibia model was acquired from the BodyParts3D database^31^. To prove potential for manufacturing customs parts for soft robotics or prosthetics, a hand model was fabricated. (**Figure 8D**). A right hand was scanned with a smart phone using the Polycam application (Altadena, CA, USA) and the hard exterior of the model was 3D printed. After printing, PhSC was injected into the molds to obtain the hand model. Moreover, to demonstrate the potential for integration of 3D circuitry and sensors into PhSC, a piezoresistive pressure sensor was utilized (**Figure 8E**). The sensor was integrated into the lower palm part of the hand model. As the pressure was applied, the resistivity changed, which was detected by a voltage drop (**Movie 3**). For future applications, PhSC can be used to produce soft robotics or custom prosthetics with integrated microfluidics and circuitry.

**Figure 8.**
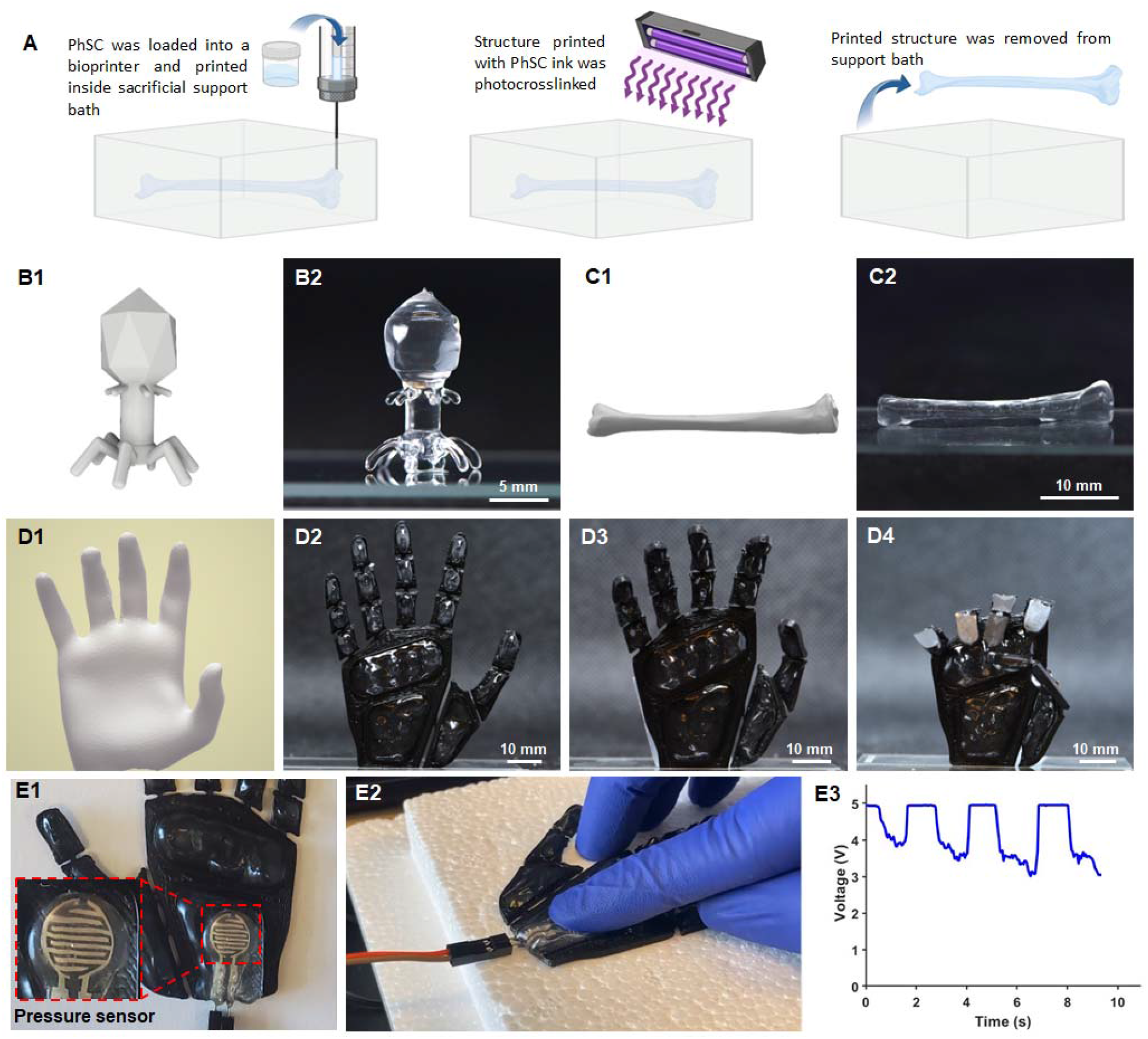
PhSC as an ink. (A) A schematic of embedded printing of the PhSC ink. Bacteriophage (B1) model and (B2) its printed form. Tibia (C1) model and (C2) its printed form. (D1) A scanned right hand and manufactured hand model in (D2) open, (D3) half-closed, and (D4) closed form. (E1) An integrated pressure sensor. (E2) Pressure sensor being pressed on. (E3) Pressure detection via voltage drop.

### 3.6. Biocompatibility of PhSC

Along with fabrication of microfluidic devices and printing of PhSC, we demonstrated its potential for cell culture work. Initially, biocompatibility of PhSC was tested with HDFs. HDFs were grown on PhSC (that were previously coated with 0.1% (w/v) gelatin) for seven days and it was found that cell viability remained higher than 90% (**Figure S5**). Biocompatibility of PhSC material was further tested with ADSCs. ADSCs were seeded onto PhSC that had 0.1% gelatin coating and cells were grown for seven days. ADSCs were stained with calcein-AM (living cells; stained green) and EthD-1 (dead cells; stained red) for the viability assessment (**Figure 9A**). After Day 7, cell viability remained higher than 90% (**Figure 9B**). Alamar Blue reduction assay demonstrated that cells continued to proliferate on PhSC surfaces for seven days (**Figure 9C**). These findings show that PhSC was biocompatible and suitable for potential biological applications, such as fabrication of PhSC-based organ-on-a-chip devices.

**Figure 9.**
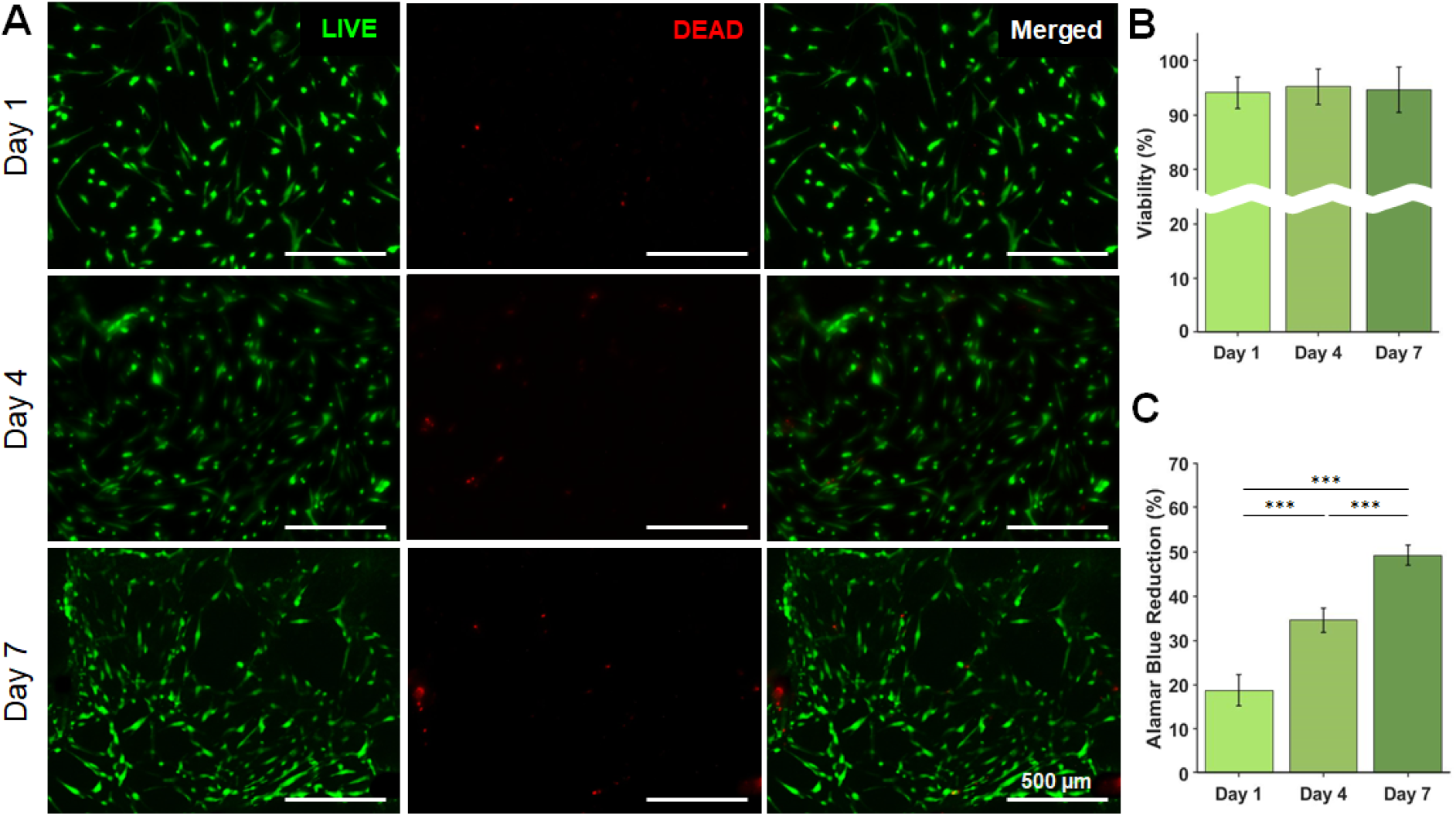
Biocompatibility of PhSC. (A) LIVE/DEAD images and (B) viability of ADSCs at Days 1, 4, and 7. (C) Alamar Blue Reduction with ADSCs at Days 1, 4, and 7 (*n*=3, *p*\*\*\*<0.001).

## 4. Conclusion

This study focuses on developing transparent photocrosslinkable silicone that can be 3D printed as well as used as a support bath for embedded printing. We demonstrated that the developed silicone composite was biocompatible and highly tunable in terms of rheological and mechanical properties that extends its applications to a wide range of microfluidics and soft robotics applications as shown here with multiple proof of concepts. Extrusion printing of PhSC as an ink or embedded printing within the PhSC support bath was highly stable and yielded complex intertwined 3D constructs with high reproducibility, where microfluidic channels could be fabricated up to the scale of tens of microns. Overall, PhSC presents a research potential to easily manufacture and integrate complex devices, such as flow-focusing devices and custom prosthetic parts with integrated electronics, thereby advancing the research in the fields of microfluidics and soft robotics.

## Acknowledgement

This research has been funded by the National Institute of Health grants R01EB034566 and U19A142733. Yasar Ozer Yilmaz acknowledges the support from The Scientific and Technological Research Council of Turkey (TUBITAK) under the BIDEB/2214-A Doctoral Scholarship Program. The authors thank Penn State Media & Technology Support Services for providing high-resolution cameras for imaging purposes. The authors are also grateful to Dr. Jiang Yang (Penn State University) for providing access to material characterization equipment.

