## Supplementary Information for "A Versatile Photocrosslinkable Silicone Composite for 3D Printing Applications"

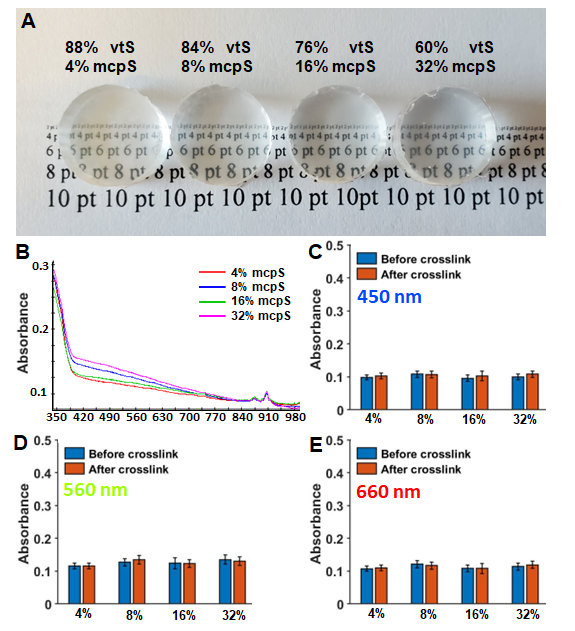

**Figure S1.** **Transparency of PhSC.** (A) Photographic presentation showing the transparency of PhSC. (B) UV-Vis spectrum of PhSC mixtures. Absorbance at (C) 450, (D) 560, and (E) 660 nm (*n=*6).

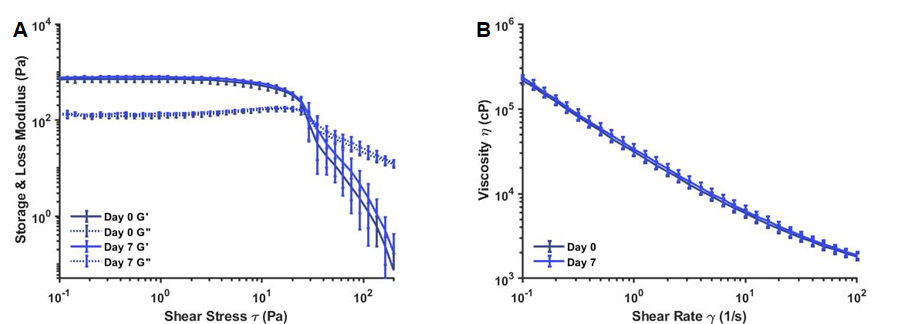

**Figure S2. Time dependency of PhSC.** Rheological properties of PhSC on Days 0 and 7 (*n=3*).

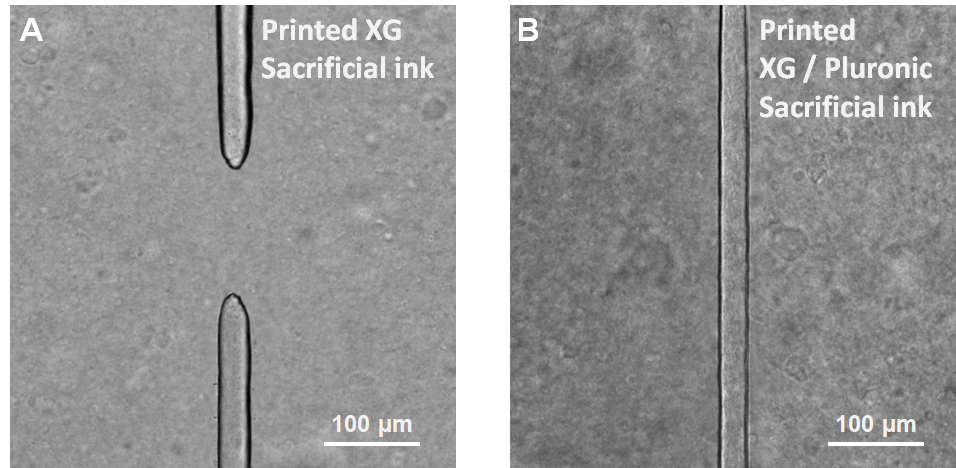

**Figure S3. Sacrificial ink selection.** (A) A failed filament with 35 µm width printed using XG as a sacrificial ink. (B) A filament with 30 µm diameter printed using XG/Pluronic as a sacrificial ink.

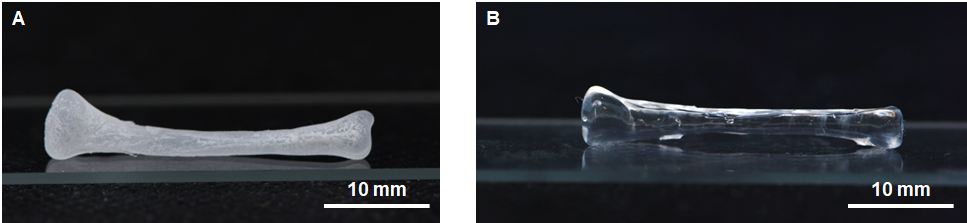

**Figure S4. Embedded printing in different support baths.** A tibia model printed with PhSC inside a (A) hydrophobic and (B) hydrophilic support bath.

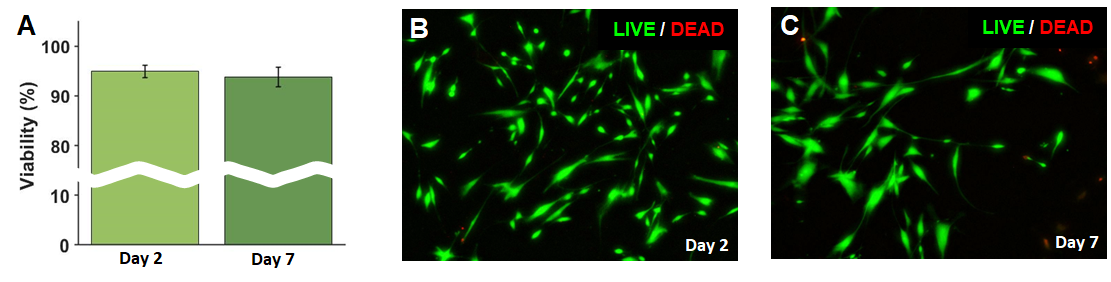

**Figure S5.** **Biocompatibility of the PhSC surface with HDFs .** (A) LIVE/DEAD assay showing the viability of HDFs. LIVE/DEAD image of of HDFs on 0.1% gelatin-coated PhSC surface on (B) Days 2 and (C) 7 (*n=*3).

**Movie Captions:**

**Movie 1: Generation of microrods in a flow focusing device**

**Movie 2: Generation of Janus microrods in a flow focusing device**

**Movie 3: Integrated pressure sensor**
